# Neural coding for tactile motion: Scanning speed or temporal frequency?

**DOI:** 10.1101/2025.09.06.674674

**Authors:** Yu-Po Cheng, Jian-Jia Huang, Chun-I Yeh, Yu-Cheng Pei

**Affiliations:** Department of Physical Medicine and Rehabilitation, Chang Gung Memorial Hospital, Linkou, Taoyuan City, Taiwan; Institute for Neuroscience, Texas A&M University, College Station, TX, USA; Master of Science Degree Program in Innovation for Smart Medicine, Chang Gung University, Taoyuan City, Taiwan; Department of Psychology, National Taiwan University, Taipei City, Taiwan; Neurobiology and Cognitive Science Center, National Taiwan University, Taipei City, Taiwan; Graduate Institute of Brain and Mind Science, College of Medicine, National Taiwan University, Taipei City, Taiwan; School of Medicine, Chang Gung University, Taoyuan City, Taiwan; Center of Vascularized Tissue Allograft, Chang Gung Memorial Hospital, Linkou, Taoyuan City, Taiwan

**Keywords:** speed, temporal frequency, tactile motion, somatosensory cortex, single unit

## Abstract

Humans effortlessly perceive the speed of an object moving across their fingers, but how the brain encodes this information, especially across the hierarchical stages in the primary somatosensory cortex, remains unclear. This study thus investigated coding schemes, including rate and temporal codes, for tactile motion speed in macaque S1 areas 3b, 1, and 2. Extracellular electrophysiology recorded single-unit activities when a rotating sinusoidal grating ball of a fixed spatial period (wavelength of 1, 2, or 4 mm) was presented on the fingerpad at various speeds (20–320 mm/s). The results showed that the rate code was commonly employed to differentiate the stimulus scanning speed, spatial period, and scanning direction across S1 regions. In contrast, the temporal code was used to faithfully represent the stimulus temporal frequency, which was defined as the speed divided by the spatial period. Notably, area 3b had a wider range of frequency responses than did areas 1 and 2. These findings demonstrate that S1 uses both rate and temporal codes to encode distinct aspects of tactile motion. Future research should investigate how temporal patterns in S1 neuronal activity are potentially transformed and utilized in downstream somatosensory areas to form tactile motion perception and guide perceptual decisions.

## Introduction

We obtain tactile information about an object by either actively moving our fingers across a surface or passively allowing the object to move against our fingerpads. Thus, humans can effortlessly perceive key motion attributes, including speed ^1,2^, direction of motion ^3–7^, and surface texture ^8–10^. Extensive research has been undertaken to understand how humans perceive tactile motion, from initial reception by mechanoreceptors and processing in the peripheral and central nervous systems to perception and behavioral responses such as psychophysical functions.

The process begins with mechanoreceptors in the skin, where rapidly adapting type 1 (RA1), slowly adapting type 1 (SA1), and Pacinian corpuscle (PC) receptors transform mechanical force into neural signals. Distinct response properties were found among these receptors: the RA1 and SA1 receptors encode lower-frequency stimuli with small receptive fields, whereas PC receptors cover higher frequencies, are often related to vibration, and have broader receptive fields ^11^. These signals are transmitted using afferent pathways to the primary somatosensory cortex (S1). Within S1, anatomical divisions were found among areas 3b, 1, and 2 ^12^, along with increasing receptive field size and complexity from area 3b to areas 1 and 2 ^13–15^, suggesting a hierarchical processing stream ^16^. Despite the insights into mechanoreceptor properties and anatomical divisions, how tactile motion information is processed and potentially transformed in S1 and subsequently downstream remains unclear, especially considering that S1 receives and integrates inputs from other sources.

To characterize tactile motion processing, psychophysical and electrophysiological studies are two complementary methods: one focuses on the behavioral outcomes as a function of stimuli, and the other focuses on the working neural mechanism. Psychophysical studies have revealed the ability of humans to discriminate motion information related to the human body, to perceive stimuli with various sensitivities, and to provide an estimate of stimulus speed ^1,4,17^. However, the perceived tactile moving speed is not simply the ratio of the perceived distance to the perceived duration of motion ^1^, unlike the calculation of veridical motion speed. Moreover, the speed estimate is influenced by the surface properties of the moving stimuli. Research using rotating patterned drums ^18–20^ and spheres ^21^ shows that human subjects report that speed estimates monotonically increase as the surface moving speed increases for both periodic and nonperiodic dot-patterned surfaces ^18^ and that rougher patterns (with a spatial period of 8 mm) could lead to slightly lower speed estimates than their smooth counterparts (2 & 3 mm). Analogously, the performance for speed discrimination differs across texture surfaces ^8^. Given that the geometric pattern of moving tactile stimuli can influence both perceived speed and neurophysiological activities ^8,22^, it is crucial to explore whether the coding schemes for speed and texture share common upstream information and the extent to which speed can be encoded regardless of spatial characteristics.

To investigate the neural basis of these perceptions, electrophysiological studies have characterized the representations of peripheral afferents and neurons in the cerebral cortex. Recording from afferent fibers in monkeys ^8,23,24^ and humans ^25,26^ revealed that SA1, RA1, and PC fibers are activated by tactile motion. The mean spiking rate monotonically increases with increasing motion speed in mechanoreceptive fibers ^26^ and neurons in primate S1 ^8,15,27–29^. This scheme of rate codes, which assumes that information is encoded by spiking rates, prevails in most of the literature. However, temporal features are robust in terms of neural response in afferents ^25^, which indicates that the scheme of temporal codes may also contribute to speed tuning.

In contrast to the rate code, the temporal code assumes that information is encoded by the precise timing and temporal patterns of spikes. The investigation of neuronal responses to stimuli’s temporal characteristics has shifted from periodic skin indentations ^30,31^ to moving stimuli contacting the skin surface ^32^. As a moving object indents and stretches the skin, its speed and surface spatial pattern determine the skin’s vibrational profile ^19,33,34^, whose fine temporal information could be crucial for perceiving moving textures ^22^. Temporal cues of tactile stimuli also play a determinant role in discriminating the stimulus spatial pattern ^35^ and moving speed ^18^. Although peripheral mechanoreceptive afferents can faithfully replicate the temporal pattern of tactile stimuli ^25,30^, there is no consensus on which coding scheme—rate, temporal, or both—S1 neurons employ to represent tactile motion speed. Therefore, we aimed to examine the coding mechanism for tactile motion speed in S1 regions representing the fingerpad and how this representation might change across the hierarchical processing stages of areas 3b, 1, and 2.

In the present study, by single-unit recording in *macaque* monkeys, we examined how S1 neurons can use **temporal codes** to encode the speed or temporal frequency of tactile motion. The presentation of sinusoid grating engraved on the rotating ball on the fingerpads offers us a unique approach to do so ^21^. The results showed that although a substantial portion of S1 neurons still encoded the speed via rate code, a majority of neurons used temporal code to encode the temporal frequency of the stimuli. Among S1 neurons, areas 3b, 1, and 2 applied different coding schemes by showing that the stimulus-evoked spiking rate peaked at 40–160 mm/s in a great portion of area 3b neurons, whereas most neurons in area 2 exhibited a monotonically increasing spiking rate with increasing speeds. Additionally, area 3b had a greater proportion of neurons representing the stimulus temporal pattern than areas 1 and 2 did.

## Results

### Tactile motion encoded by the rate code

To determine how S1 neurons encode tactile motion information, we performed extracellular electrophysiology in three *macaque* monkeys (Supplementary Material and Supplementary Table 1). We applied rotating stimulus balls with spatial periods of 1, 2, and 4 mm to the fingerpad at speeds of 20, 40, 80, 160, and 320 mm/s (Figure 1A−C). For each single unit, we calculated the stimulus-evoked spiking rates (Figure 1G & H) and periodicity metrics of the spike trains (Figure 1I & J), including the autocorrelogram with added noise (see Methods) and its power spectrum. Out of 195 recorded units, 158 were responsive to tactile motion (area 3b, n=55; area 1, n=58; area 2, n=45), based on their spiking rates (sample single unit shown in Figure 1G−J). We applied a three-way ANOVA (direction × speed × spatial period) to categorize the feature selectivities of these units. As a result, we identified 33/55, 35/58, and 34/45 direction-selective; 40/55, 48/58, and 39/45 speed-selective; and 54/55, 52/58, and 39/45 spatial period-selective single units from areas 3b, 1, and 2, respectively. No significant difference was found in the proportions of direction-selective units (area 3b: 60.0%, area 1: 60.3%, area 2: 75.6%; *X*^2^ = 3.31, *p* = 0.191, chi-square test), speed-selective units (area 3b: 72.7%, area 1: 82.8%, area 2: 86.7%; *X*^2^ = 3.35, *p* = 0.187), or spatial-period-selective units (area 3b: 94.5%, area 1: 89.7%, area 2: 86.7%; *X*^2^ = 1.83, *p* = 0.400) among the S1 regions (*Supplementary Table 2*). In addition, no regional difference was identified for single units that showed interaction effects (*Supplementary Table 2*).

**Figure 1.**
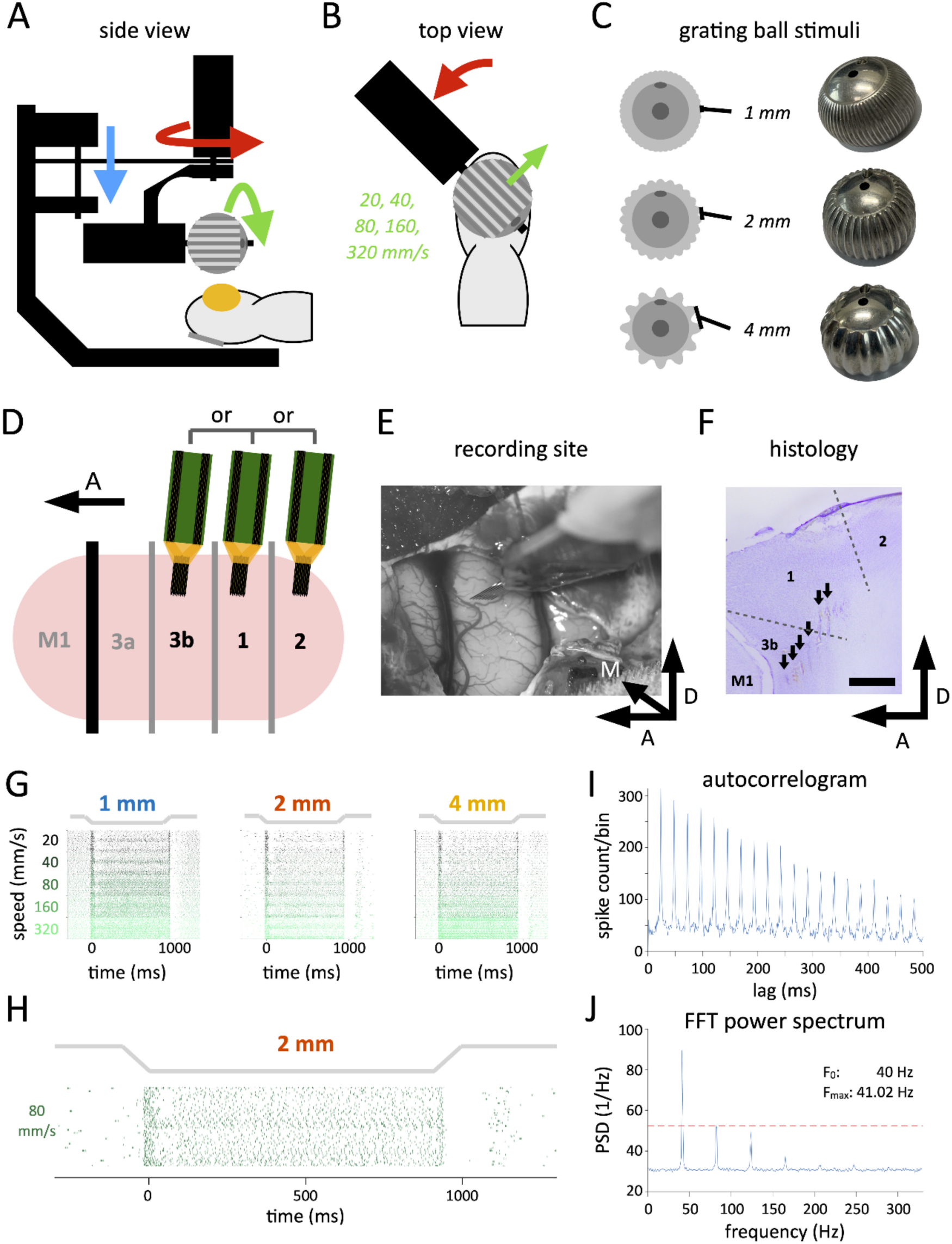
Experimental setup and assessment of the periodicity of neural activities in a sample single unit. (A) Side view of the tactile stimulator setup. A miniature tactile stimulator with a sinusoid grating ball and three motors, each of which was used to present the scanning orientation (red), scanning speed and direction (green), and indentation (blue), was used. Tactile motion stimuli were delivered by the stimulator to the animal’s fingerpad (yellow). (B) Top view of the tactile stimulator setup. The scanning speed was 20, 40, 80, 160, or 320 mm/s. (C) Axial view of the grating balls. Three sinusoidal grating balls with spatial periods of 1, 2, or 4 mm were used. (D) The recording areas of interest. Areas 3b, 1, and 2 in the primary somatosensory cortex (S1) were recorded. The black line indicates the central sulcus, and the gray lines indicate the boundaries between brain areas in S1. A: anterior. (E) Sixty-four-channel silicon-based electrode lowered perpendicularly into the recording site. A, anterior; M, medial; D, dorsal. (F) An example recording site shown on the post hoc histology for S1. The trajectories of the probe electrode are observed in the Nissl-stained brain slice. Electrocaudation at certain channels of the recording electrode was performed to verify the precise location of the recordings. The black arrows indicate the recording electrode tracts. A, anterior; D, dorsal. Scale bar =1 mm. (G) Raster plots under all spatiotemporal stimulus conditions recorded from a sample single unit. The black-to-green color gradient indicates scanning speeds ranging from 20 to 320 mm/s. The spatial period of the scanning ball (blue: 1 mm, orange: 2 mm, yellow: 4 mm) is shown from the left to right panels. (H) Raster plot of the sample unit’s neural activity in response to 2-mm grating ball scanning at 80 mm/s (black line). Time 0 indicates the time the stimulus ball fully indents the fingerpad. (I) Periodicity of the autocorrelogram of neural responses in Panel H, showing a series of peaks with a fixed interval of ∼ 24 ms. Their amplitude gradually decreases with time. (J) Power spectral density (PSD) of the sample autocorrelogram in Panel I. The red dashed line indicates the mean + 5×SD, a criterion used to detect significant frequency components. Only the frequency component of the maximum power met the criteria.

### Temporal code preserved information about stimulus temporal frequency

The question remains whether S1 neurons encode the stimulus, temporal frequency, or both, particularly through the temporal code. We hypothesized that if S1 neurons encode the stimulus speed, the responses would be linearly correlated with speed regardless of the spatial period. Conversely, if S1 neurons encode the stimulus temporal frequency (stimulus F_0_), the responses would instead correlate with temporal frequency (see the Methods). The temporal patterns of spike trains from a sample single unit were analyzed through power spectra in response to combinations of stimulus speeds and spatial periods (Figure 2A) and from the prestimulus baseline (Figure 2B). The results indicated that the **frequency of maximum power** (F_max_, see the Methods & Figure 1G−J) was positively correlated with the stimulus scanning speed, although a constant offset was observed between spatial periods (Figure 2C). Importantly, F_max_ (perfectly) aligned with stimulus F_0_ (Figure 2D), suggesting that the spiking pattern of a single unit may follow a temporal frequency coding scheme.

**Figure 2.**
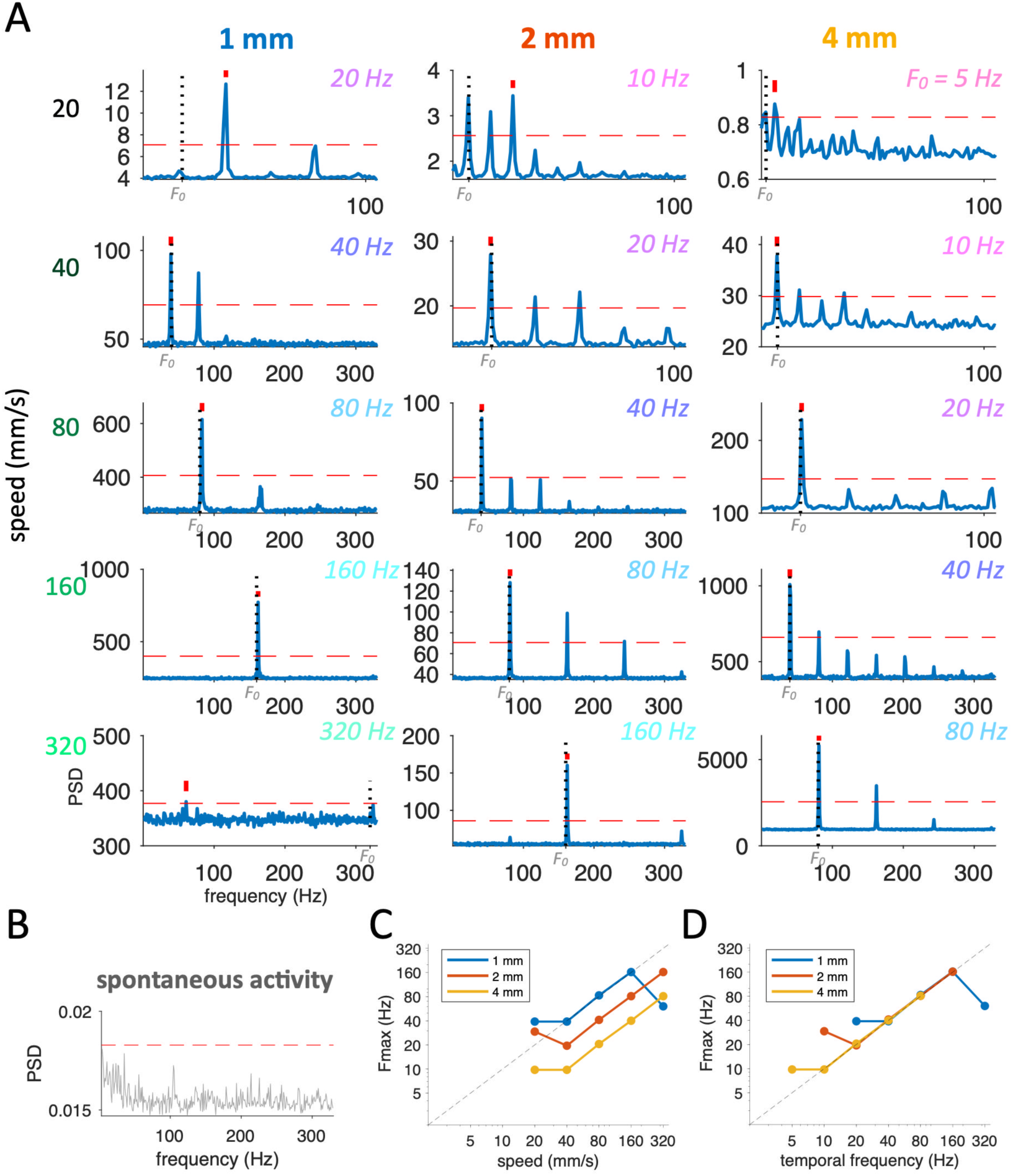
Power spectra and PSD of the sample single unit for the spatiotemporal combinations. (A) Power spectra of the sample singe unit in Figure 1G for the spatial period and different stimulus speed combinations. The red bars indicate the frequency of maximum power (F_max_) among the observed frequencies, the red dashed lines indicate the criteria for significant frequency components, and the black dashed lines indicate stimulus F_0_. The row panels are displayed by stimulus speed, and the column panels are displayed by spatial period. The temporal frequency is labeled using a color patch with a pink-to-cyan gradient ranging from 5 to 320 Hz. (B) Average PSD of spontaneous neural activity before stimulus onset (across trials). (C) Correspondence between F_max_ and stimulus speed. The offset of the curves of the three spatial periods was observed, with the longer spatial period having a lower F_max_. The diagonal dashed line indicates the condition when F_max_ is identical to the scanning speed. (D) Correspondence between F_max_ and the temporal frequency of the stimulus. The sample unit showed a F_max_ that was nearly identical to stimulus F_0_ across spatial periods in the 10–160 Hz range. In contrast to the offset of the curves of the three spatial periods as observed in C, the diagonal dashed line indicates the condition when F_max_ is identical to stimulus F_0_.

Next, we considered two possibilities for S1 neurons: (1) a temporal frequency coding scheme (**TF coding**) or (2) a speed coding scheme (**SP coding**). Given our experimental design, these schemes would yield distinct response patterns regarding F_max_ versus stimulus F_0_. The TF coding scheme predicts that F_max_ is proportional to the scanning speed and inversely proportional to the spatial period (Figure 3A left); in contrast, the SP coding scheme predicts that F_max_ is proportional to the scanning speed but independent of the spatial period (Figure 3A right). Additionally, we defined neurons that follow either coding scheme only **for a subset** of the tested spatial periods as SP-like coding or TF-like coding (e.g., F_max_ aligns with stimulus F_0_ for spatial periods of 2 and 4 mm or for 4 mm alone).

**Figure 3.**
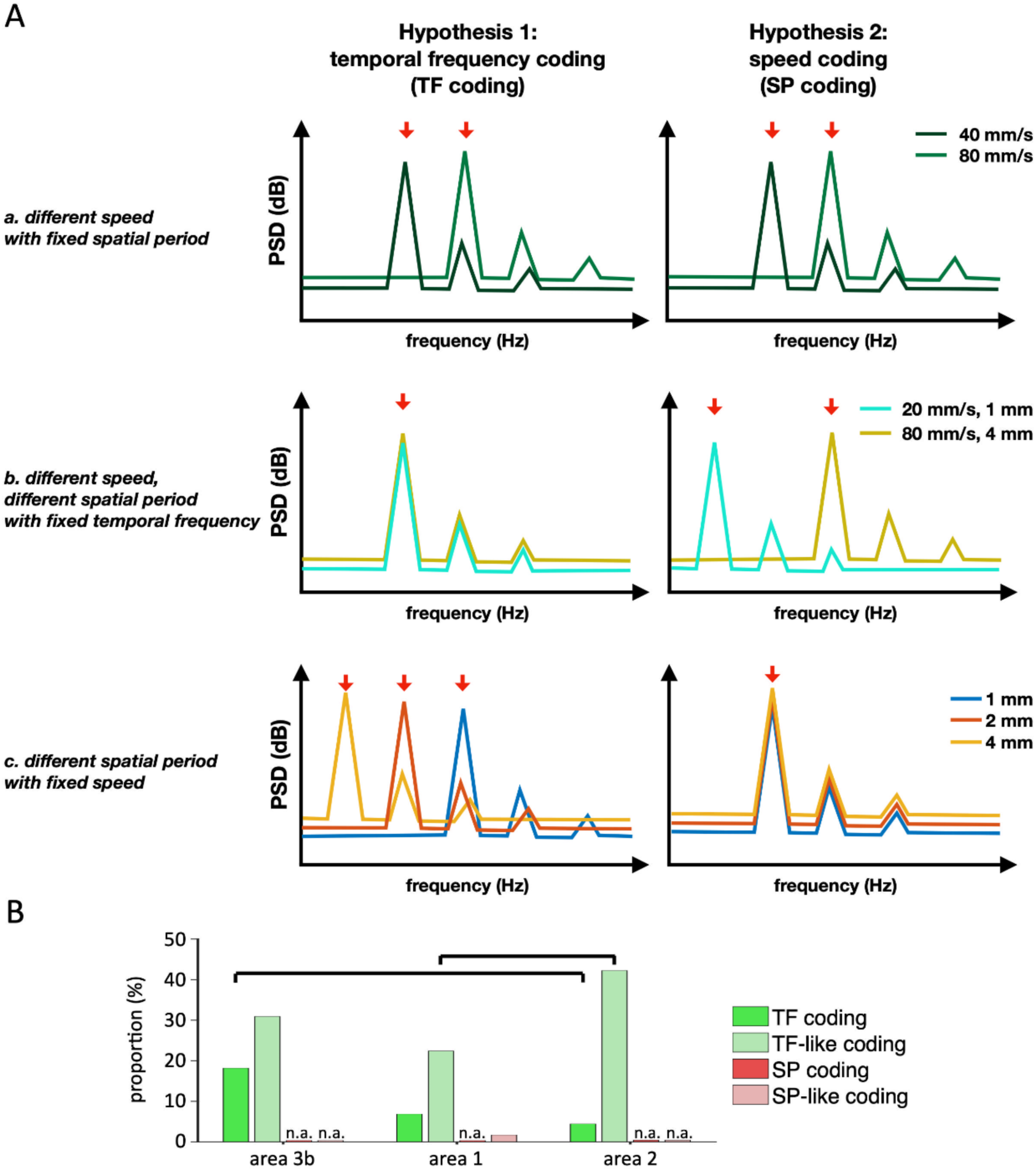
Two hypotheses proposed using the presentation of PSD in the frequency domain for identifying putative SP (speed) and TF (temporal frequency) coding schemes. (A) In the left column, we hypothesize that if a neuron encodes the stimulus using the TF coding scheme, the neuron is tuned to the temporal frequency, which is defined as the speed divided by the spatial period. Thus, F_max_ is proportional to the scanning speed and inversely proportional to the spatial period. The expected F_max_ would (a) increase with increasing speed, (b) remain the same for combinations of speed and spatial period, resulting in a fixed temporal frequency, and (c) decrease with increasing spatial period. In the right column, we hypothesize that if a neuron encodes the stimulus using the SP coding scheme, its F_max_ will be proportional to the scanning speed and independent of the spatial period. The expected F_max_ would (a) increase with increasing speeds for a fixed spatial period, (b) increase with increasing speeds for changing spatial periods, and (c) be independent of the change in the spatial period at a fixed speed. (B) The proportions are defined as the number of single units of each coding scheme divided by the number of single units in each area. Each unit is analyzed for the degree to which the neuron’s tuning is compatible with each coding scheme in each of the 15 spatiotemporal conditions. The results showed that a majority of single units follow the TF and TF-like coding schemes in all three areas. The black lines indicate statistical significance (*p* < 0.05, chi-square test).

**Figure 4.**
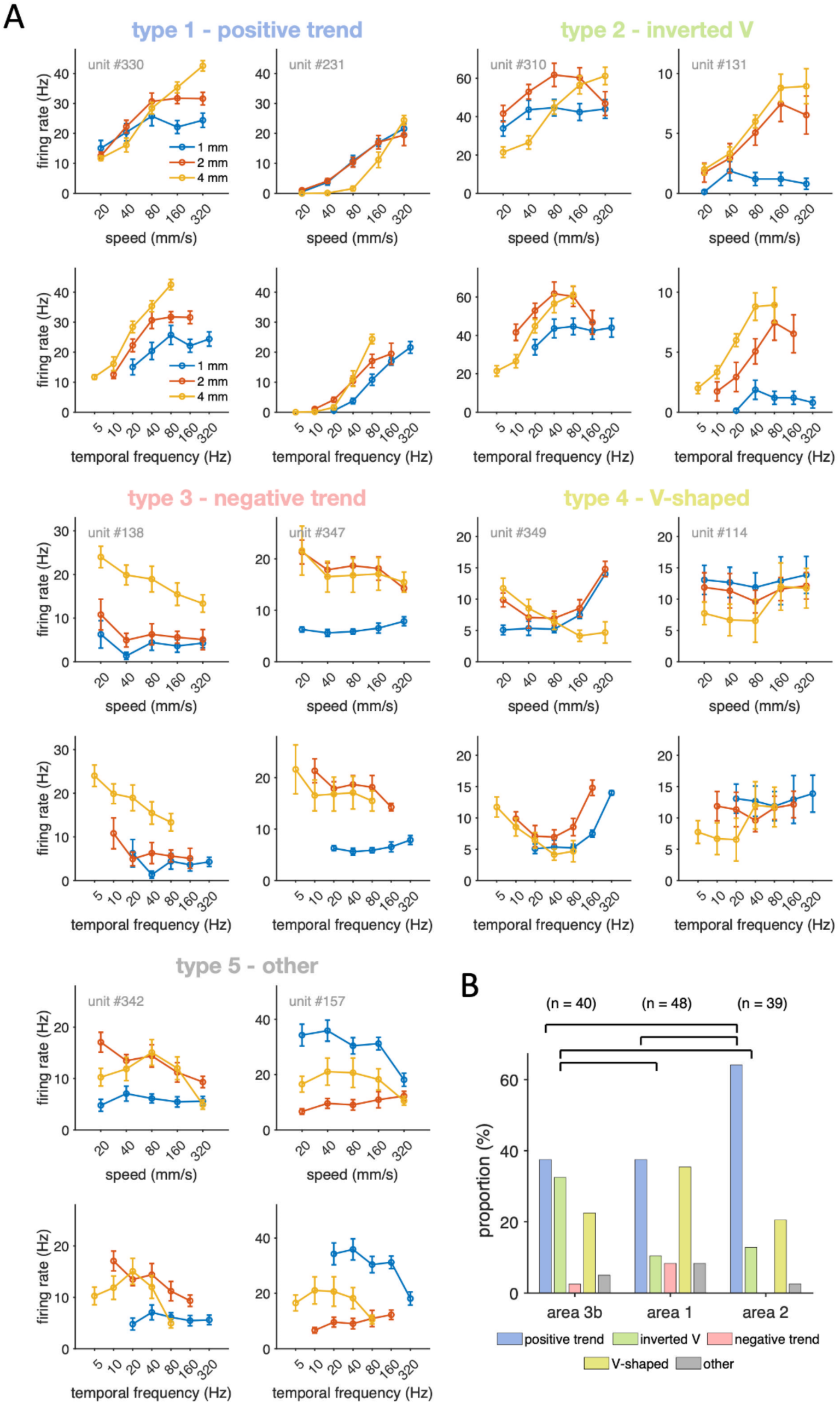
Five types of speed selectivity and effects of temporal frequency, spatial period, and speed on neural spiking rates. (A) The speed-selective units were categorized into one of the five types based on their firing rates as a function of stimulus speed. The five speed-selectivity types are the positive trend, inverted V, negative trend, V-shaped, and other patterns, and two sample units are illustrated. The top panels show the firing rate as a function of stimulus speed, and the bottom panels show the firing rates as a function of temporal frequency. The blue, orange, and yellow curves indicate conditions with spatial periods of 1, 2, and 4 mm, respectively. (B) Proportion of single units in the five speed-responsive types and their distribution among regions of S1. The majority are positive trend units in area 3b (38%), area 1 (38%), and area 2 (64%). The black lines indicate statistical significance (*p* < 0.05, chi-square test). Error bars: s.e.m.

Regional differences among areas 3b, 1, and 2 revealed that 18% of the neurons in area 3b followed TF coding, which was significantly greater than the 4% in area 2 (*p* = 0.037, chi-square test) but did not differ from the 7% in area 1 (*p* = 0.071, chi-square test). This finding indicates the invariance of TF coding across spatial periods (Figure 3B). Approximately 31% of the neurons in area 3b demonstrated TF-like coding, which may be related to the lower evoked spiking for a spatial period of 1 mm, as shown in Figure 4A (unit #330). In area 1, 7% of the neurons followed TF coding, with 22% showing TF-like coding. In area 2, only 4% of the neurons followed TF coding, whereas 42% of the neurons presented TF-like coding, a significantly greater proportion than that in area 1 (*p* = 0.032, chi-square test). These findings suggest that neurons in area 2 extract specific tactile features rather than general spatiotemporal properties. Notably, no units in area 3b, 1, or 2 exhibited SP coding, and only 1.72% of the neurons in area 1 followed SP-like coding.

In summary, TF coding schemes emerged as the primary coding mechanism in S1. Neurons in area 3b can encode a wide range of temporal frequencies, whereas neurons in areas 1 and 2 are more specialized, responding primarily to specific combinations of scanning speeds and spatial periods.

### Characterizing speed-selective responses

To further examine the contribution of the rate code, we categorized the speed-selective units into five types: positive trend, inverted V, negative trend, V-shaped, and other patterns (see the Methods for details, Figure 4A). Most S1 units showed a positive correlation between speed and spiking rate, particularly within the positive trend and inverted V types. Notably, the proportion of positive trend units was significantly greater in area 2 (64%) than in areas 3b (38%; *p* = 0.018, chi-square test) and 1 (38%; *p* = 0.014, chi-square test). The proportion of inverted V-type units was significantly greater in area 3b (33%) than in areas 1 (10%; *p* = 0.011, chi-square test) and 2 (13%; *p* = 0.038, chi-square test). The negative trend units constituted only 3% of area 3b, 8% of area 1, and 0% of area 2. No regional differences were found for the V-shaped type

(area 3b: 21%, area 1: 35%, area 2: 21%) or for the other patterns (area 3b: 5%, area 1: 8%, and area 2: 3%) (Figure 4B). Most units of the inverted V type had peak spiking rates of 160 mm/s in areas 3b, 1, and 2. Moreover, the V-shaped type had the lowest spiking rates of 80 mm/s (area 3b), 40 mm/s (area 1), and 40 mm/s (area 2) (*Supplementary* Figures 5*−7*).

Additionally, we examined whether the spiking rate of S1 neurons was mediated by speed or temporal frequency, with temporal frequency defined as speed divided by the spatial period of the stimulus ball. When the temporal frequency was used as the x-axis, we observed nearly identical slopes for the positive trend, inverted V, and V-shaped types in area 3b at spatial periods of 2 and 4 mm (*Supplementary* Figure 5), indicating spiking activity invariant to spatial periods. In contrast, neurons in area 1 (*Supplementary* Figure 6) presented invariant spiking rates when the spatial periods were 1 and 2 mm when plotted against temporal frequency. In area 2 (*Supplementary* Figure 7), the functions of temporal frequency versus spiking rate overlapped for spatial periods of 2 and 4 mm, similar to area 3b for the positive trend and inverted V types.

We identified a regional difference for the rate code across S1, with area 3b having the greatest proportion of units tuned to 160 mm/s, whereas area 2 had the greatest proportion of units showing a monotonically increasing spiking rate with increasing scanning speeds up to 320 mm/s. Spatial period-invariant responses were identified by comparing the spiking rate with the temporal frequency instead of the scanning speed.

### Temporal frequency coding and harmonics

Several S1 neurons showed prominent peaks in the periodograms outside stimulus F_0_. Previous research has reported that the surface texture of a moving object induces skin vibrations at specific temporal frequencies and their harmonics ^19^. On this basis, we hypothesized that S1 neurons encode not only stimulus F_0_ but also vibrational information from the harmonics, where the harmonics are defined as multiples of stimulus F_0_ (Figure 5A).

**Figure 5.**
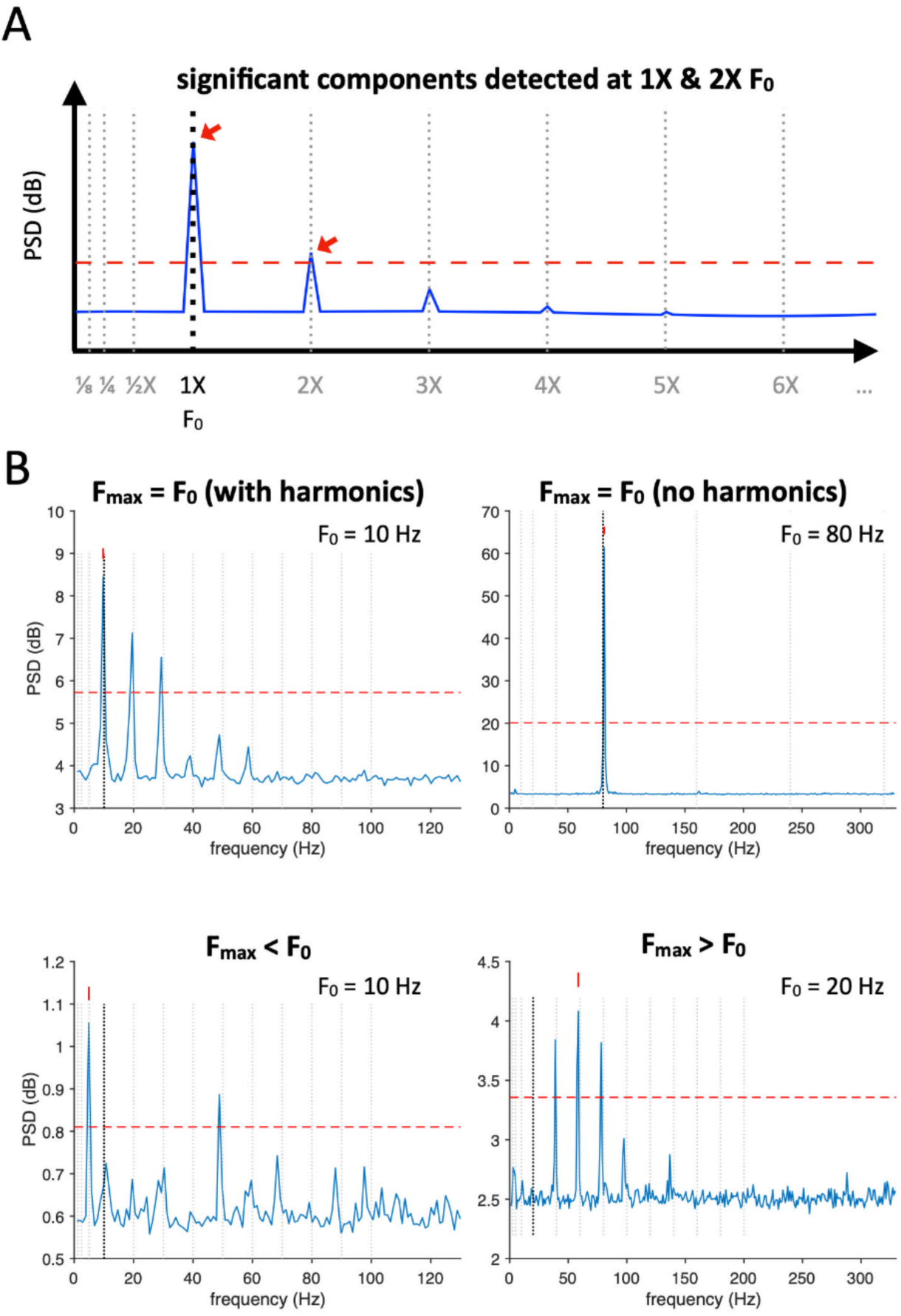
Patterns of temporal information coding in spiking responses with respect to stimulus F_0_ and its fractions or multiples. (A) Identification of significant frequency components of autocorrelations was performed after the procedure in Figure 1J. We examined whether these components fit the multiples or fractions (from 1/8× to 10×) of stimulus F_0_, defined as the harmonics. The frequency components whose peak PSD is greater than the threshold of the mean + 5×SD criterion (red dashed line) are considered significant. In this example, two peaks, 1× and 2×, of stimulus F_0_ met this criterion. (B) Four samples of neuronal spiking patterns. Top left, 1×, 2×, and 3× of stimulus F_0_ are significant, and F_max_ = stimulus F_0_. Top right, only 1× of stimulus F_0_ is significant, and F_max_ = stimulus F_0_. At the bottom left, ½× and 5× of stimulus F_0_ are significant, and F_max_ = ½× stimulus F_0_. Bottom right, 2×, 3× and 4× of stimulus F_0_ are significant, and F_max_ = 3× stimulus F_0_.

Three distinct patterns were observed among the units studied. The first group included 42% of area 3b, 17% of area 1, and 18% of area 2 and had an F_max_ equal to stimulus F_0_, along with harmonics at attenuated amplitudes in at least one of the fifteen tested conditions (5 scanning speeds × 3 spatial periods) (top left panel, Figure 5B). The second group consisted of 58% of area 3b, 48% of area 1, and 49% of area 2, which exhibited only a single frequency component (F_max_) (top right panel, Figure 5B). The third group comprised 65% of area 3b, 62% of area 1, and 60% of area 2, whose F_max_ did not match stimulus F_0_ but was a fraction or multiple of stimulus F_0_ (bottom panels, Figure 5B). Notably, all units exhibited response patterns beyond these three types in at least one of the fifteen tested conditions, indicating that S1 neurons are unable to encode the entire temporal frequency range from 5 to 320 Hz.

Our findings indicated that S1 neurons can faithfully encode stimulus F_0_ up to 160 Hz, as well as the harmonics up to five times stimulus F_0_ or more (the sample unit shown in Figure 6A). A substantial proportion of these units focused on the lower frequencies (summarized in Figure 6B). In area 3b, the proportions of units responding to stimulus F_0_ were **45% at 5 Hz, 26% at 10 Hz, 38% at 20 Hz, and 24% at 40 Hz.** A smaller proportion encoded the harmonics (e.g., 2× stimulus F_0_, **15% at 5 Hz, 24% at 10 Hz, 15% at 20 Hz, and 7% at 40 Hz**). In area 1, even fewer units (2× stimulus F_0_, less than 10% for all F_0_) exhibited spectral peak components at the harmonic frequencies, indicating a reduced capacity to temporally encode vibrational information from the harmonics. In area 2, the distribution of neurons encoding stimulus F_0_ and the harmonics was similar to that observed in area 3b but with a lower number of units.

**Figure 6.**
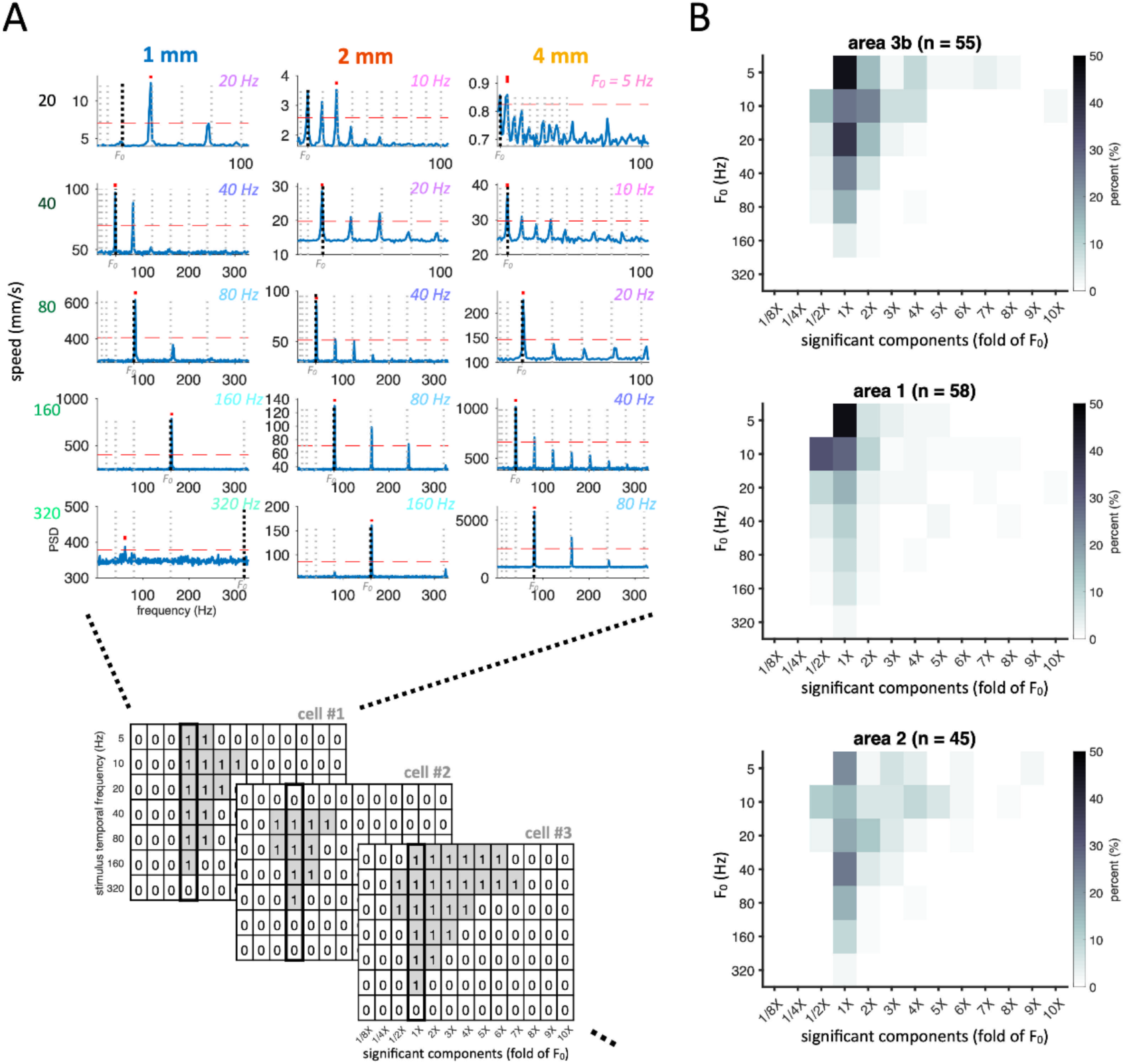
Proportions of harmonics that fit the multiples or fractions of stimulus F_0_. (A) The distribution of significant frequency components of autocorrelations among all tested conditions is summarized for the sample cell from Figure 3A (cell #1 in the illustration) and all the others. The summary table shows the successful detection of significant harmonics as 1 and failure as 0 at seven temporal frequencies (from 5 to 320 Hz) and their multiples and fractions (from 1/8× to 10×) for each cell. Summary tables from the same S1 region were accumulated to compute the regional proportion. (B) Proportion of single units that show a significant frequency response in the power spectrum to each of the multiples and fractions. At multiples of the stimuli temporal frequency (1/8×−10×), harmonic peaks were widely observed in area 3b units at low frequencies and stopped at approximately 80 Hz (4th harmonic of 20 Hz & 2nd harmonic of 40 Hz). In contrast, harmonic peaks were barely observed in area 1 units. In area 2, similar to area 3b but different, harmonic peaks were observed up to 120 Hz (3rd harmonic of 40 Hz). The different patterns in temporal frequency coding indicated distinct processing complexity among S1 regions.

In summary, our findings highlight distinct patterns of temporal coding among neurons and across S1 regions, with area 3b demonstrating the most effective encoding of detailed temporal information about the stimulus.

## Discussions

### Representation of tactile motion in the primary somatosensory cortex (S1) using rate coding or temporal coding

Peripheral mechanoreceptors encode physical properties of tactile stimuli mainly using the **rate code**. Specifically, neurons have been shown to have varying spiking rates to represent moving speed ^18,28,36^, moving direction ^37^ and surface texture ^8,19^. However, the nervous system also adopts a **temporal code**, through which neurons fire with intervals related to the stimulus’s temporal pattern ^22,25,31^. Previous studies have highlighted S1’s role in the temporal code ^8,38,39^; however, the fidelity and mechanisms of how S1 represents the temporal details of tactile motion remain unclear. The present study uses sinusoid gratings engraved on rotating stimulus balls with varying spatial periods and scanning speeds to examine how S1 neurons encode the **speed** and **spatial frequency** of tactile stimuli. Consistent with previous reports, we found that a large proportion of S1 neurons across areas 3b, 1, and 2 demonstrate varied spiking rates as a function of the direction, speed, and spatial period of tactile stimuli. Regarding the temporal patterns in neuronal spiking activities, our results indicate that most S1 neurons mainly employ temporal codes to represent the temporal frequency of stimuli while using rate codes to represent the speed of tactile motion.

Building on previous research, we further explored the role of different S1 regions in encoding tactile information through analyzing neuronal spiking activity patterns. Specifically, area 3b encodes a variety of stimulus temporal frequencies, whereas areas 1 and 2 represent narrower ranges. Salinas *et al.* ^38^ reported that periodic mechanical vibrations can evoke spike trains in S1 neurons with reliable periodicity, a pattern less frequently observed in the secondary somatosensory cortex (S2). Furthermore, some S2 neurons presented a greater spiking rate in response to low-frequency stimuli, which contrasts with the observed trend in most mechanoreceptors and S1 neurons. Additionally, the information conveyed via temporal codes was significantly greater in S1 area 3b than in area 1, whereas that in S2 approached zero. In the periphery, Weber *et al.* ^22^ reported that RA1 and Pacinian afferents might use their spiking temporal pattern to convey tactile information for discriminating the scanning texture. Similarly, our study revealed a prominent periodicity of S1 activities that correlated with the spatiotemporal pattern of tactile stimuli by mirroring the temporal pattern within a frequency range from 5 to 160 Hz. These results, together with existing literature, provide strong evidence that S1 encodes the spatiotemporal patterns of tactile stimuli through both rate and temporal codes ^39^.

### Harmonic frequencies in S1 activity

Intriguingly, S1 neuron spiking events occurred not only at fixed time intervals corresponding to the stimulus frequency but also at intervals that were fractions or multiples of it. As a result, the periodogram of S1 neurons exhibited spectral power peaks at the stimulus frequency and several of its harmonics. In some cases, the spectral power at these harmonics was equal to or exceeded that at the stimulus frequency. The neural mechanisms for this phenomenon remain unclear but could be related to the skin’s vibration profile or misalignment of input timings. Our findings suggested that S1 could represent the periodicity in the skin vibratory profile, which results from relative movements of the skin and objects, depending on the surface spatial patterns of those objects ^19,33,34^. Distinct harmonic patterns in neuronal spiking for varying surface textures may enable the nervous system to identify tactile stimuli based on past experiences. Downstream neurons (i.e., S2 and beyond) could use this information for further processing. According to the findings of Romo et al. ^40^, S2 neurons may employ rate codes for behaviorally relevant information. Thus, we speculate that the temporal profile of spiking patterns, including the harmonic feature for each surface texture, could be interpreted by neurons in downstream areas and integrated into a rate code for making behavioral decisions ^41^.

### Neural codes for speed vs. temporal frequency

Our experimental design allowed us to investigate whether S1 neurons encode stimulus speed, temporal frequency, or both, with some combinations of speed and spatial period sharing the same temporal frequency. We demonstrated that the index of spiking periodicity, F_max_, monotonically increased with increasing stimulus temporal frequency (speed/spatial period) rather than speed itself. Notably, the coding range for these frequencies varied across S1 areas, with the highest proportion of neurons for wide and narrow ranges observed in areas 3b and 2, respectively. Additionally, our analysis of neuronal spiking revealed invariant spiking rates for stimulus temporal frequency across spatial periods, further suggesting neural codes for temporal frequency. Distinct patterns were observed across the S1 region, where areas 3b and 2 showed invariance between spatial periods of 2 mm and 4 mm, whereas area 1 displayed invariance between 1 mm and 2 mm.

These findings suggest the existence of regional differences in processing complexity, potentially caused by the increasing complexity along the processing hierarchy in S1 ^16^. In alignment with human psychophysical findings on tactile speed perception ^28^, we propose that S1 neurons may prioritize the encoding of stimulus temporal frequency over speed, where surface geometric patterns such as dots and edges contribute to the dissociation of these features.

## Conclusion

This study highlights that, in addition to employing the rate code, macaque S1 neurons also encode the temporal characteristics of tactile motion through specific patterns of spiking activity. Distinct temporal code patterns were identified across areas 3b, 1, and 2. Future studies should aim to characterize the neural mechanisms underpinning the “responsive range,” referring to the range of stimulus temporal frequencies in which S1 neurons can faithfully represent, as well as further explore how the brain processes the complex harmonic content of tactile stimuli along the somatosensory pathway.

## Materials and Methods

### Experimental Animals

The studies involved three Taiwanese macaque monkeys (*Macaca cyclopis)*, two males and one female, weighing between 5 and 8 kg. All experimental procedures were conducted in accordance with the National Institutes of Health Guide for the Care and Use of Animals. The animal care and handling procedures were followed, and the protocols were approved by the Institutional Animal Care and Use Committee of National Yang-Ming University (IACUC: 1040714).

### Study Design

Tactile stimuli were delivered using a miniature tactile stimulator (Figure 1A−B), which was developed specifically for presenting motion stimuli at various speeds and in various directions ^21,42^. The spatial pattern of these stimuli was determined by engravings on the stimulus balls ^43^, allowing for the application of gratings with different spatial periods. The miniature stimulator consists of three individual motors, each of which performs (1) rotation to control the speed of motion, (2) orientation to control the direction of motion, and (3) vertical excursion to control the depth of indentation onto the skin. This design allows for precise control of the speed, direction, indentation depth, and wavelength of the scanning sinusoid grating.

Three aluminum sinusoid grating balls with a diameter of 20 mm and a trough-to-peak ridge height of 1 mm were used (Figure 1C). These stimulus balls had spatial periods of 1, 2, or 4 mm. The animals’ forearms were placed palm-up, and each of the fingers being stimulated was fixed using a finger holder. The tactile motion was presented by one of the three sinusoidal gratings, with the grating ball indented into the fingerpad at a depth of 1 mm during stimulation. Each of the grating balls was tested in a separate session, with rests interleaved between sessions. In each session, the grating ball was presented in various directions ranging from 0° to 315° in 45° increments, and at different surface scanning speeds of 20, 40, 80, 160, or 320 mm/s, in pseudorandom order. The coordinates of the motion directions were somatotopic, with 0° and 90° indicating the rightward and distal directions, respectively.

Each trial consisted of a stimulus period lasting 1 second, followed by an interstimulus interval of 1 second. The whole experiment had 3 sessions (one for each ball with different spatial periods) × 8 directions × 5 speeds, yielding a total of 120 combinations, each repeated 10 times. The stimulus temporal frequency for each testing condition was calculated using the following equation:

Temporal frequency = speed/spatial period.

### Surgical Preparation & Electrophysiology

The animals were initially anesthetized with ketamine hydrochloride (10 mg·kg^-1^ i.m.) and then maintained under neurolept anesthesia (20–70 mcg·kg^-1^·h^-1^ i.v. fentanyl and 0.3–0.5% isoflurane) and a skeletal muscle relaxant (1.2 mcg·kg^-1^·h^-1^ i.v. rocuronium bromide) throughout the experiment. Anesthesia was continued with 1.5–3.5% isoflurane while the animals were artificially ventilated with 5% end-tidal CO_2_. To ensure that the anesthesia remained at stage two, an electroencephalographic wire was implanted under the skull. Toe pinch testing was regularly performed to detect any pain responses for further monitoring of the status of anesthesia. Vital signs were closely monitored using an electrocardiogram, a pulse oximeter, and a rectal thermometer, to maintain heart rates of 120–180 beats per minute, S_P_O_2_ ≥ 98%, and a rectal temperature of approximately 38 °C.

A rectangular craniotomy was performed overlying S1 according to Horsley−Clarke coordinates (anterior 6, lateral 21 mm), with a width of 15 mm. We first manually mapped the brain regions within the craniotomy window by inserting multiple tungsten microelectrodes (NAN Electrode Drive System, Nazareth Illit, Israel) to identify the receptive fields in Brodmann areas 1, 2, and 3b, as well as their corresponding finger areas (Chen, Friedman, Ramsden, LaMotte, & Roe, 2001; Pei, Denchev, Hsiao, Craig, & Bensmaia, 2009), by step indention of a rod or sweeping a Q-tip. Following durotomy, silicon-based multishank electrodes (NeuroNexus, A8x8-5mm-200-200-413-A64), a type of multielectrode array (MEA), were inserted to obtain signals from areas 1, 2, and 3b (Figure 1D, E). The MEA was oriented orthogonally with respect to the central sulcus to ensure that the receptive fields of the neurons were predominantly located at the same finger, a property that was then verified by manual mapping.

### Spike Sorting & Data Processing

Spike sorting for single units was performed using offline sorting (Plexon, TX, USA), through which the spike waveform and resting spiking rates across time were monitored to validate the consistency of recordings from the same single unit. Specifically, waveform consistency was verified using Pearson correlation across any two recording periods, through which a confusion matrix of correlation coefficients was constructed for every single unit, and the criterion of a waveform-consistent single unit was a coefficient ≥ 0.8. In addition, the stability of spontaneous spiking was analyzed by confirming that there were no significant changes in the spontaneous rate between the 2-second time window before and after the entire stimulation period using the Wilcoxon rank-sum test.

For every single unit, spike timing was binned using a 2-ms bin and aligned to the stimulus onset, yielding a raster plot and a perievent time histogram (PSTH). The resting, on, sustain, and off periods were defined as −200 to −100 ms, 50 to 150 ms, 200 to 300 ms, and 1000 to 1100 ms after stimulus onset, respectively, which is defined as the putative time at which the rotating ball contacts the fingertip. Single units with a mean spiking rate < 5 Hz in all of the resting, on, sustain, and off periods were excluded from further analysis.

Single units showing a significant main effect of scanning speed on the spiking rate during the sustain period using ANOVA were considered speed-selective. These units were then categorized into one of the following 5 speed-responsive types according to their spiking rates as a function of stimulus speed: (1) positive trend, with the lowest spiking rate at 20 mm/s and the highest at 320 mm/s; (2) inverted V, with the lowest spiking rate at 20 mm/s and the highest at 40, 80, or 160 mm/s; (3) negative trend, with the lowest spiking rate at 320 mm/s and the highest at 20 mm/s; (4) V-shaped, with the lowest spiking rate at 40, 80, or 160 mm/s; and (5) other patterns if the units did not fit any of the aforementioned patterns (Figure 4A).

### Periodicity analysis for stimulus-driven neuronal responses

To investigate the type of tactile motion information encoded using **temporal coding**, we determined the frequency of maximum power (F_max_) and its harmonics from the neuronal responses. The following procedures were used to obtain representative power spectra for stimulus-driven neuronal responses.

First, autocorrelograms of the PSTHs were constructed, and linear detrending was applied (Figure 1G−I). Notably, variant spiking rates can induce bias in subsequent frequency−domain analyses derived from this autocorrelogram. To address this, we implemented a **noise addition preprocessing** step ^44,45^. Specifically, normal white noise with a mean of 0 and standard deviation of the maximum peak amplitude in the autocorrelogram was superimposed on the autocorrelogram.

The resulting autocorrelgrams were analyzed using fast Fourier transform (FFT). The aforementioned noise addition preprocessing step was performed for 5000 iterations, and the resultant power spectrum among iterations was averaged to yield the final power spectrum (Figure 1J). We then searched the **frequency of maximum power (F_max_)** in the final power spectrum inside the frequency range of interest. The periodicity analysis focused on the region of interest of the temporal code of a frequency range of 3 to 330 Hz.

### Determination of coding schemes

We defined a single unit as following the speed coding scheme when its F_max_ is positively correlated with the stimulus scanning speed and is independent of spatial periodicity. Moreover, a single unit follows the temporal frequency coding scheme when its F_max_ matches stimulus F_0_. A tolerance range of +-3 Hz centered at stimulus F_0_ was used to detect the coding scheme.

### Statistics

The responsiveness to the tactile stimulus was estimated by comparing the stimulus-driven spiking rates in the on, sustain, and off periods ^46^ to the spiking rate of the resting period using the Wilcoxon rank-sum test (*p* < 0.017, Bonferroni adjustment). Three-way ANOVA was used to identify the main effects of direction, speed, and spatial period on the stimulus-driven spiking rates of individual units. A single unit was classified as direction-selective if it showed a main effect of scanning direction (*p* < 0.01) ^28^. The same criteria were applied to the main effects of speed, spatial period and all interaction terms. Chi-square tests were used to compare the proportions of responsive, direction-selective, speed-selective, spatial-period-selective, and interaction effects across S1 regions.

## Supplementary materials

We isolated 205 recorded high-quality units from the Brodmann areas 3b (n = 73), 1 (n = 78) and 2 (n = 54) of three macaque monkeys (*Supplementary Table 1*) for analysis. The stimulus-responsive units (67/73 in area 3b, 77/78 in area 1, and 51/54 in area 2) were characterized by on, off, and sustain periods. We hypothesized that the on response encodes the initial contact, the off response encodes the indentation step, and the sustain response encodes most of the information, such as the scanning speed, stimulus texture, and temporal frequency of contacts. Therefore, we focused on 158/205 units (55 in area 3b, 58 in area 1, and 45 in area 2) with significant differences in spiking rates between the sustain periods and the prestimulus baseline to investigate how S1 represents tactile speed.

## Author contributions

Conceptualization, J.J.H. and Y.C.P.; methodology, Y.P.C., J.J.H., C.I Y. and Y.C.P.; software, Y.P.C., J.J.H. and Y.C.P.; validation, Y.P.C., J.J.H. and Y.C.P.; formal analysis, Y.P.C., J.J.H. and Y.C.P.; investigation, Y.P.C., J.J.H., C.I Y. and Y.C.P.; resources, J.J.H., C.I Y. and Y.C.P.; writing—original draft preparation, Y.P.C., J.J.H. and Y.C.P.; writing—review and editing, Y.P.C., J.J.H., C.I Y. and Y.C.P.; visualization, Y.P.C., J.J.H. and Y.C.P.; supervision, J.J.H. and Y.C.P.; project administration, J.J.H. and Y.C.P.; funding acquisition, J.J.H., C.I Y. and Y.C.P. All authors have read and agreed to the published version of the manuscript.

## Acknowledgements

We thank the following funding sources:

NHRI-EX101-10113EC

NSC 100-2321-B-182-013

MOST-108-2320-B-002-030

MOST-109-2320-B-002-014

CMRPG3M1651-2 for data analysis

---

**Supplementary Figure 1.**
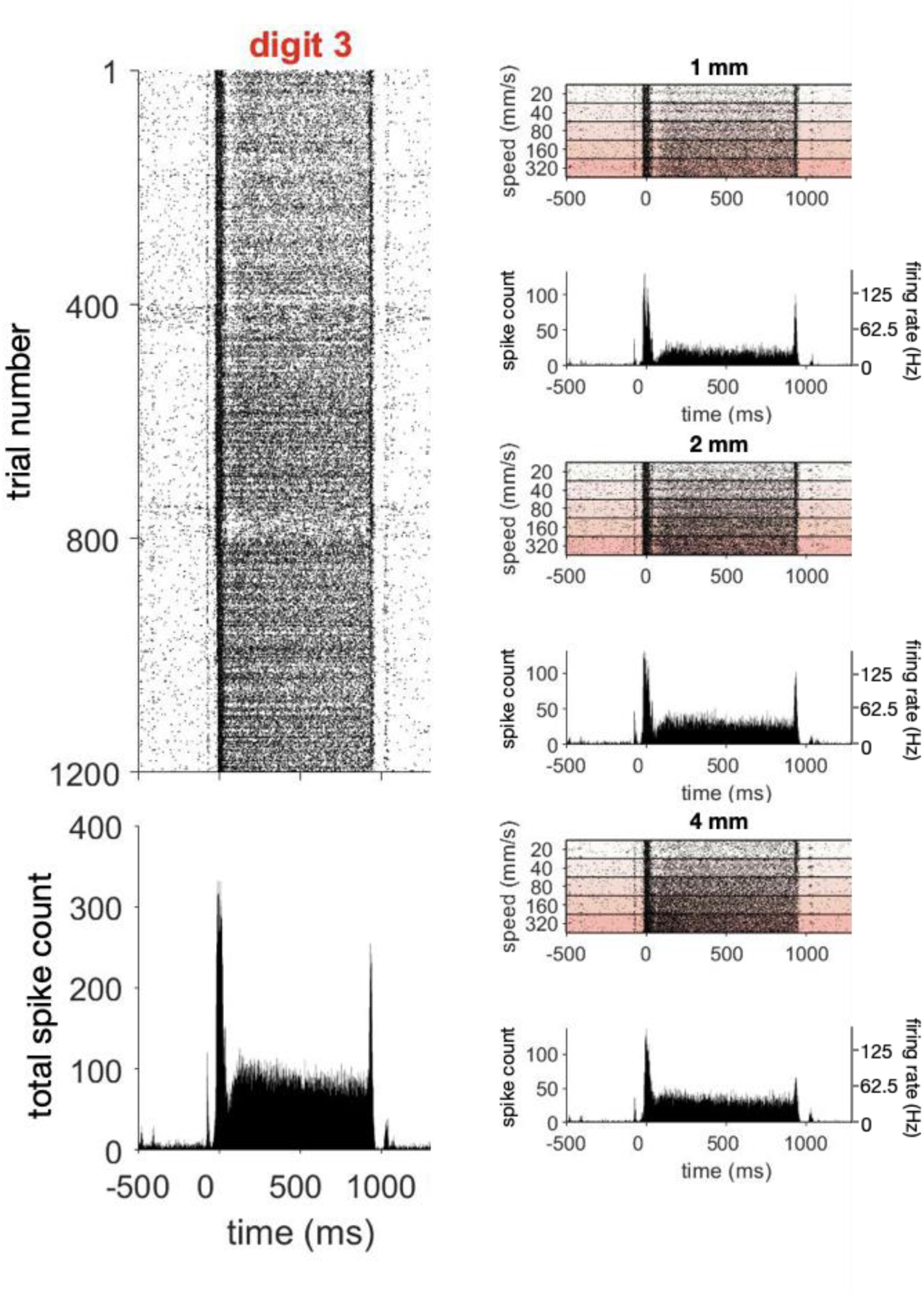
**Sample unit in area 3b.** Left column, raster plots and PSTHs of the sample unit in area 3b among all the testing trials. Time 0 indicates that the stimulus ball was fully intended in the fingerpad. Right column, breakdown raster plots and PSTHs of the sample unit based on the stimulus parameters (top panel: spatial period of 1 mm, middle panel: 2 mm, bottom panel: 4 mm; from low to high scanning speeds).

**Supplementary Figure 2.**
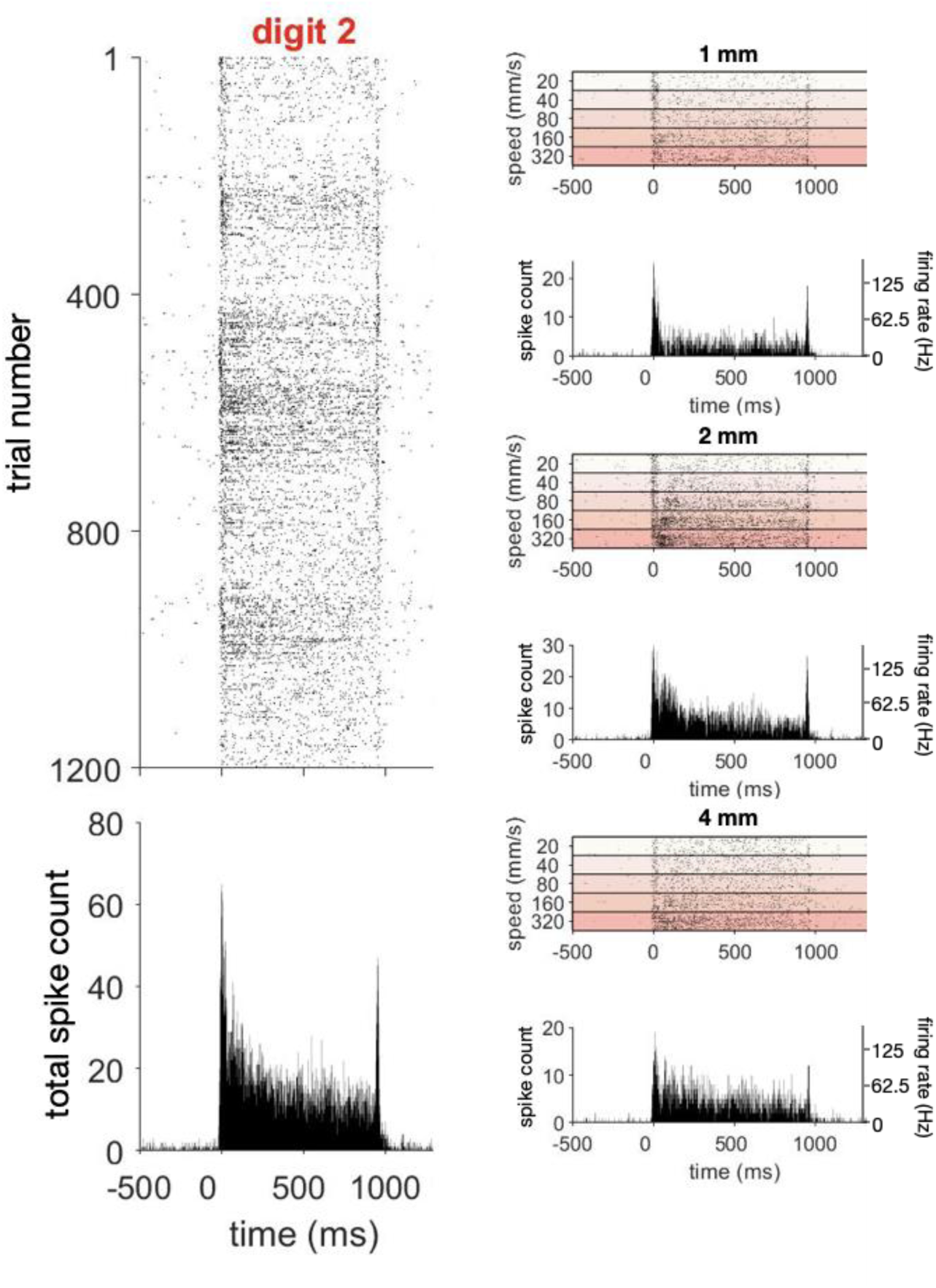
**Sample unit in area 2.** Left column, raster plots and PSTHs of the sample unit in area 2 among all the testing trials. Time 0 indicates that the stimulus ball was fully indented in the fingerpad. Right column, breakdown raster plots and PSTHs of the sample unit based on the stimulus parameters (top panel: spatial period of 1 mm, middle panel: 2 mm, bottom panel: 4 mm; from low to high scanning speeds).

**Supplementary Figure 3.**
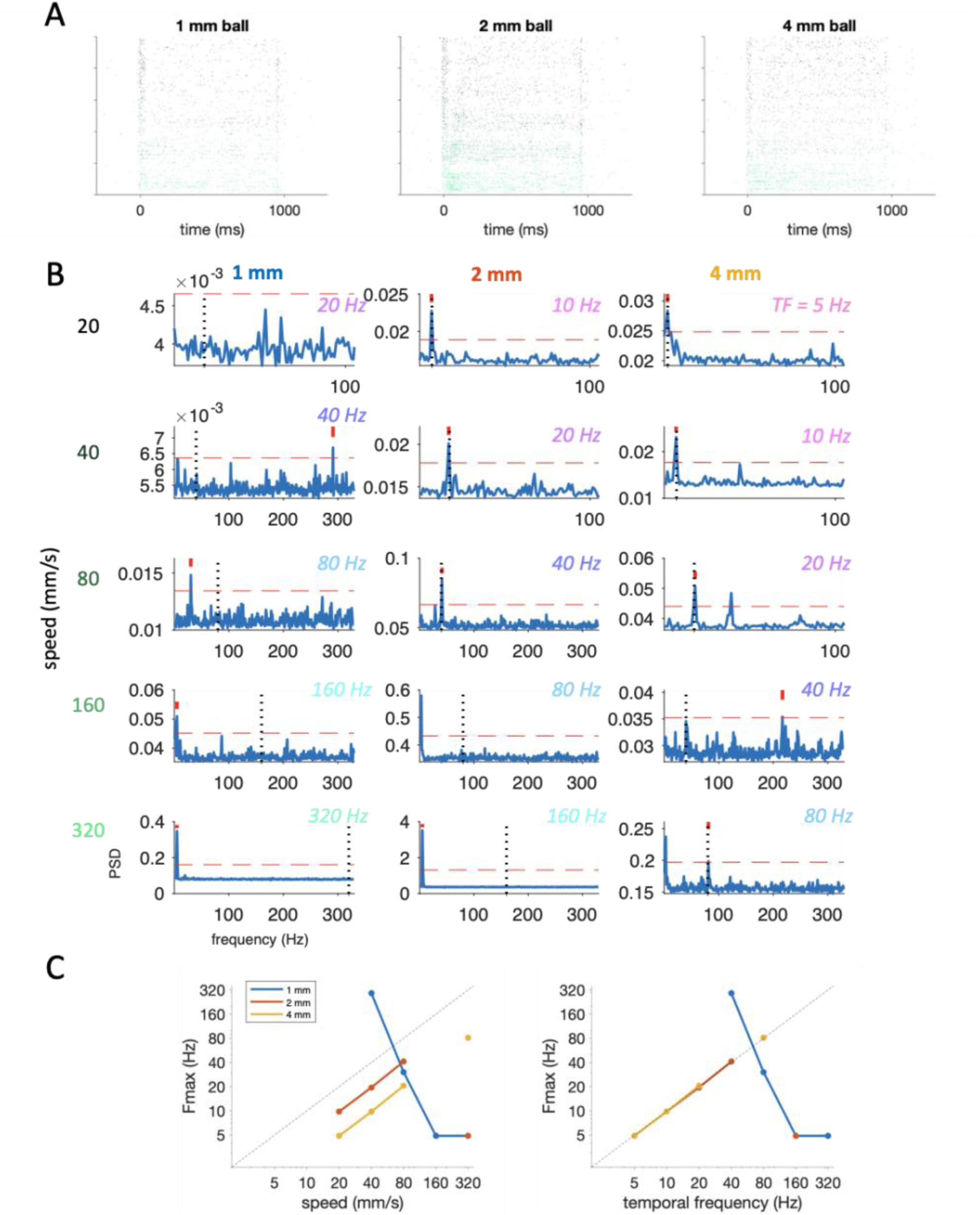
**Sample unit for the TF-like coding scheme.** (A) Raster plots of the sample single units for all spatiotemporal combinations. The black-to-green color gradient indicates scanning speeds from 20 to 320 mm/s. The spatial periods of the scanning balls (blue: 1 mm, orange: 2 mm, and yellow: 4 mm) are shown in the panels from left to right. (B) Power spectra of a single unit in A for these combinations. The red vertical lines indicate the frequency of maximum power (F_max_) among the observed frequencies; the red dashed lines indicate the criteria for meaningful frequency components; the diagonal dashed line indicates the condition when F_max_ is identical to the stimulus temporal frequency (F_0_). The row of PSD panels represents the scanning speed, and the column represents the spatial period. The temporal frequency, defined as the scanning speed over the spatial period, was labeled using a color patch with a pink-to-cyan gradient (from 5 to 320 Hz). (C) Correspondence between F_max_ and stimulus speed. The spatial period of the stimulus ball is indicated by blue (1 mm), orange (2 mm), and yellow (4 mm) lines. The sample single unit had a negative slope for the spatial period of 1 mm and two positive, comparable slopes for the spatial periods of 2 and 4 mm. An offset was observed between the lines for the 2 mm and 4 mm conditions. The diagonal dashed line indicates the condition when F_max_ is identical to the scanning speed. (D) Correspondence between F_max_ and the temporal frequency of the stimulus. The two lines for the 2 mm and 4 mm conditions overlapped the diagonal line. The diagonal dashed line indicates the condition when F_max_ is identical to stimulus F_0_.

**Supplementary Figure 4.**
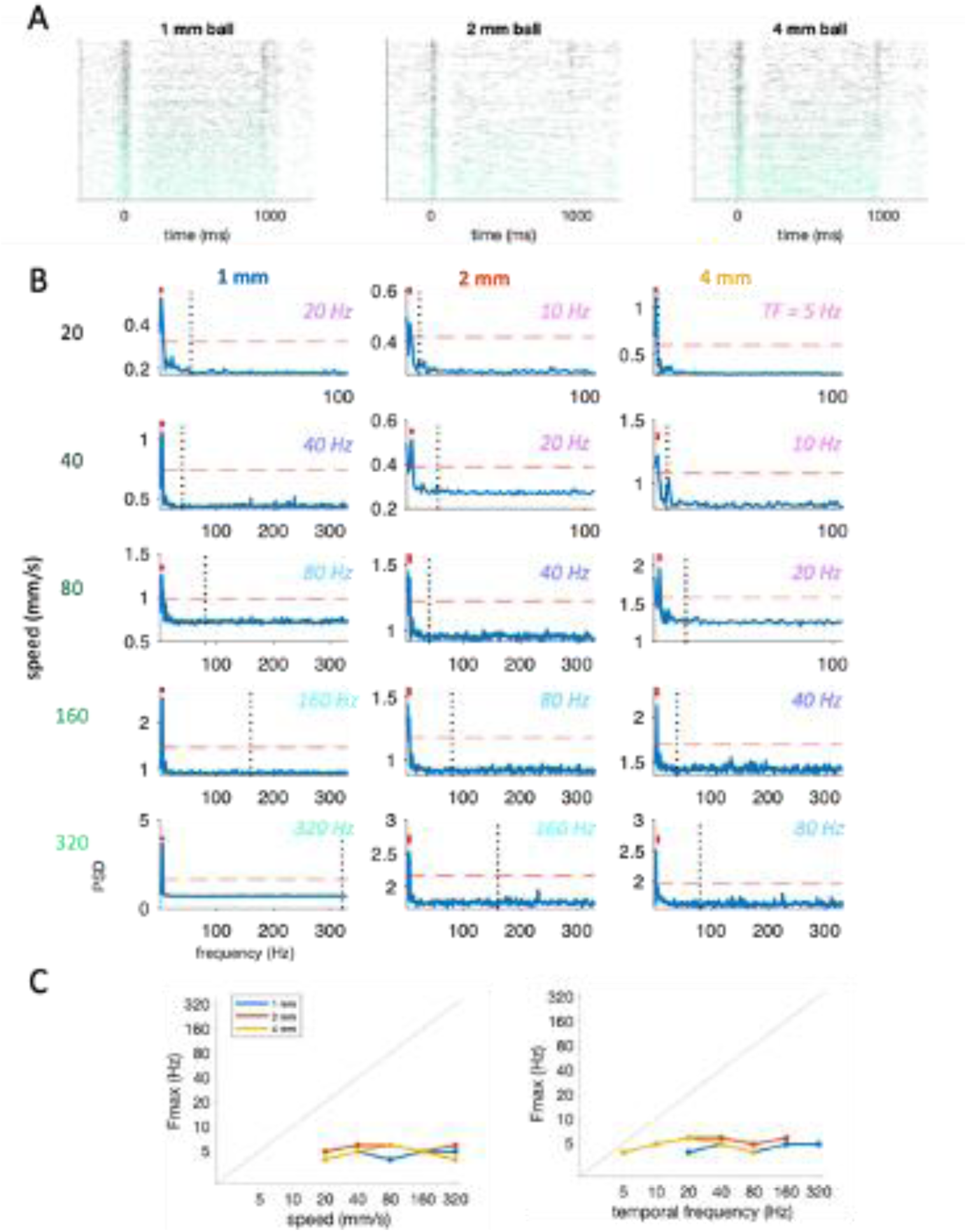
**Sample unit for the SP-like coding scheme.** (A) Raster plots of the sample single units for all spatiotemporal combinations. The black-to-green color gradient indicates scanning speeds from 20 to 320 mm/s. The spatial period of the scanning ball (blue: 1 mm, orange: 2 mm, and yellow: 4 mm) is shown in the panel from left to right. (B) Power spectra of a single unit in A for these combinations. The red vertical lines indicate the frequency of maximum power (F_max_) among the observed frequencies; the red dashed lines indicate the criteria for meaningful frequency components; and the diagonal dashed line indicates the condition when F_max_ is identical to the stimulus temporal frequency (F_0_). The row of PSD panels represents the scanning speed, and the column represents the spatial period. The temporal frequency, defined as the scanning speed over the spatial period, was labeled using a color patch with a pink-to-cyan gradient (from 5 to 320 Hz). (C) Correspondence between F_max_ and stimulus speed. The spatial period of the stimulus ball is indicated by blue (1 mm), orange (2 mm), and yellow (4 mm) lines. For the 2-mm and 4-mm spatial periods, the sample single unit showed comparable F_max_ values among spatial periods at certain scanning speeds (20–80 mm/s) and increased F_max_ with increasing scanning speed. The diagonal dashed line indicates the condition when F_max_ is identical to the scanning speed. (D) Correspondence between F_max_ and the temporal frequency of the stimulus. No specific pattern was observed for the F_max_ distribution over temporal frequency among the spatial periods. The diagonal dashed line indicates the condition when F_max_ is identical to stimulus F_0_.

**Supplementary Figure 5.**
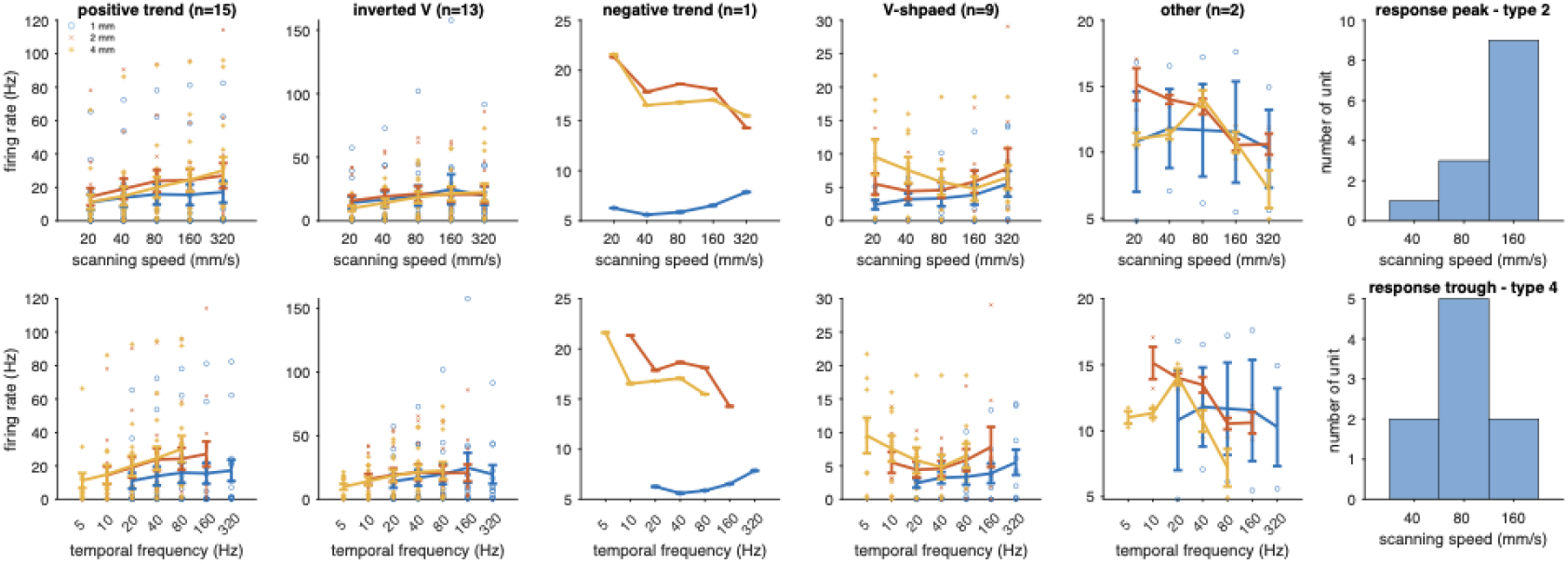
**Five types of speed selectivity in area 3b.** Means and s.e.m. values for all single units of the same type. The top panels show spiking rates as a function of scanning speed, and the bottom panels show spiking rates as a function of temporal frequency. The blue, orange, and yellow dots indicate the trials for spatial periods of 1, 2, and 4 mm, respectively. For types 2 (inverted V) and 4 (V-shaped), histograms of the response peak or trough show the scanning speeds at which the single units are tuned.

**Supplementary Figure 6.**
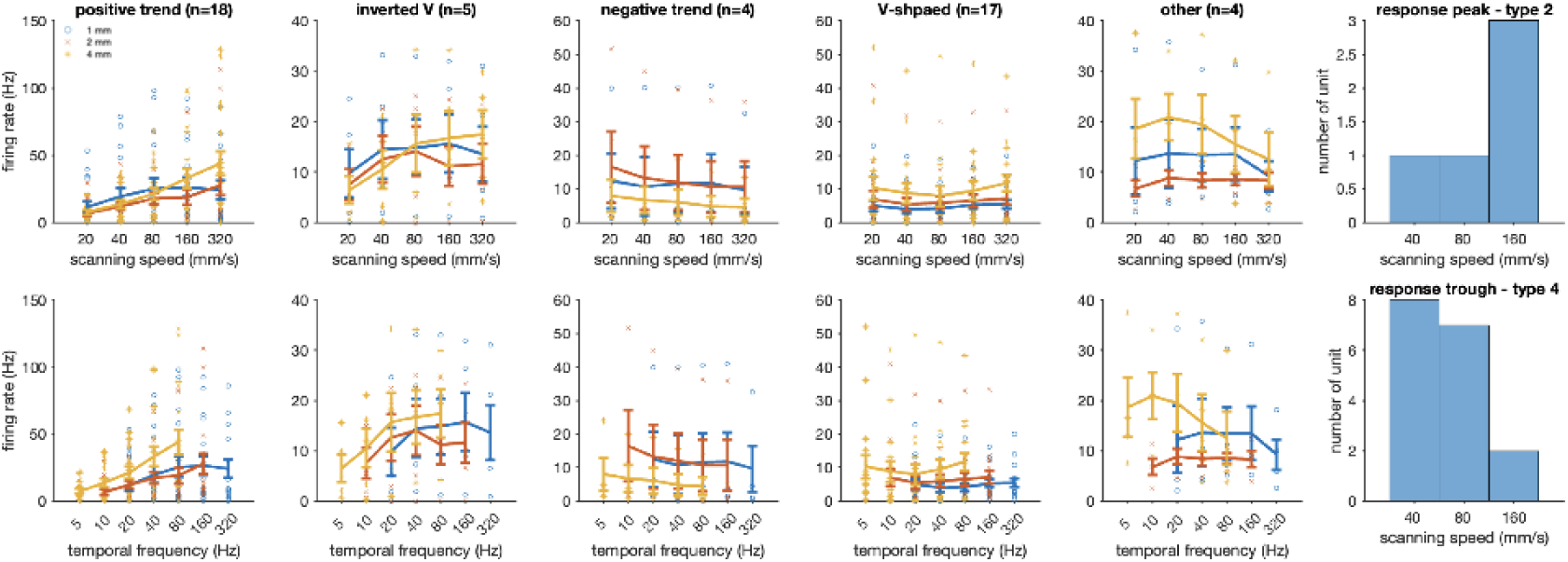
**Five types of speed selectivity in area 1.** Means and s.e.m. values for all single units of the same type. The top panels show spiking rates as a function of scanning speed, and the bottom panels show spiking rates as a function of temporal frequency. The blue, orange, and yellow dots indicate the trials for spatial periods of 1, 2, and 4 mm, respectively. For types 2 (inverted V) and 4 (V-shaped), histograms of the response peak or trough show the scanning speeds at which the single units are tuned.

**Supplementary Figure 7.**
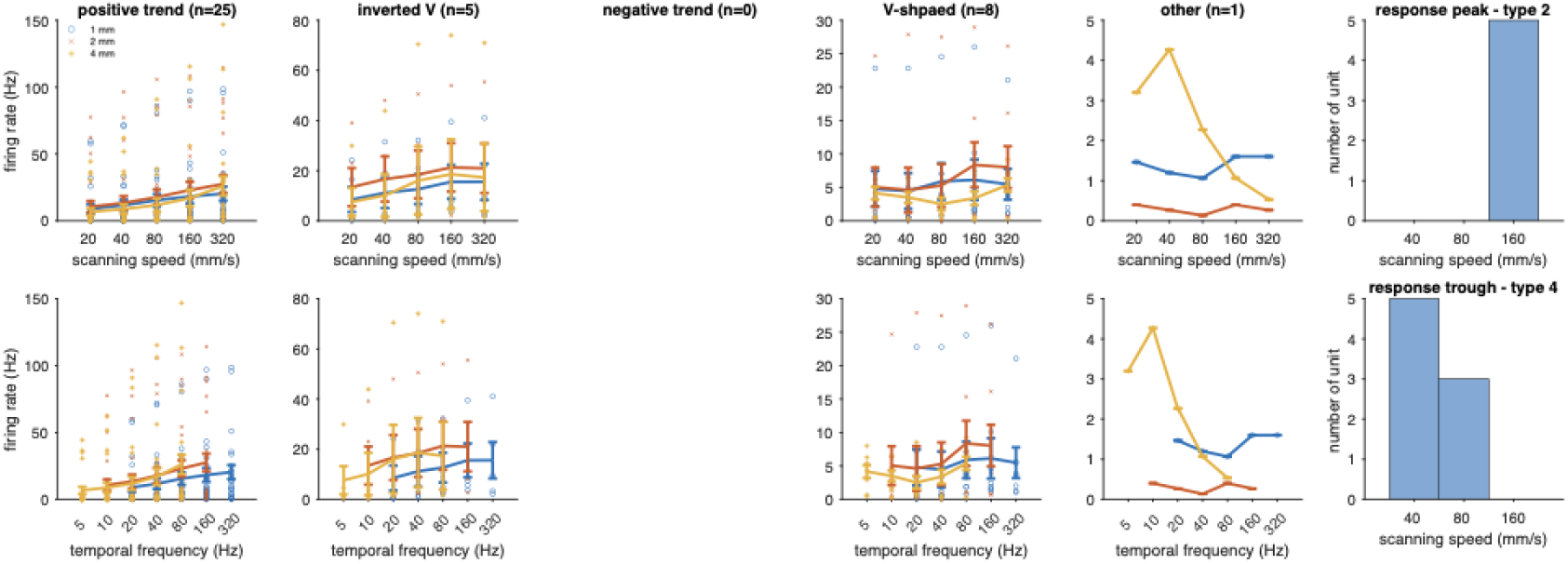
**Five types of speed selectivity in area 2.** Means and s.e.m. values for all single units of the same type. The top panels show spiking rates as a function of scanning speed, and the bottom panels show spiking rates as a function of temporal frequency. The blue, orange, and yellow dots indicate the trials for spatial periods of 1, 2, and 4 mm, respectively. For types 2 (inverted V) and 4 (V-shaped), histograms of the response peak or trough show the scanning speeds at which the single units are tuned.

**Supplementary Table 1.**
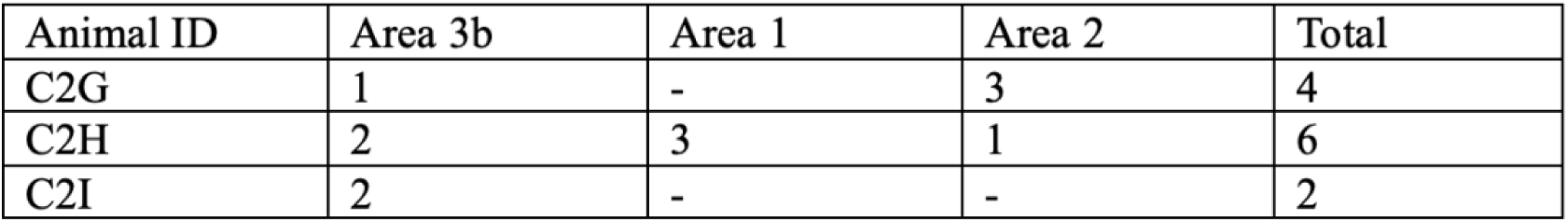
Recording sites.

| Animal ID | Area 3b | Area 1 | Area 2 | Total |
| --- | --- | --- | --- | --- |
| C2G | 1 | - | 3 | 4 |
| C2H | 2 | 3 | 1 | 6 |
| C2I | 2 | - | - | 2 |

**Supplementary Table 2.**
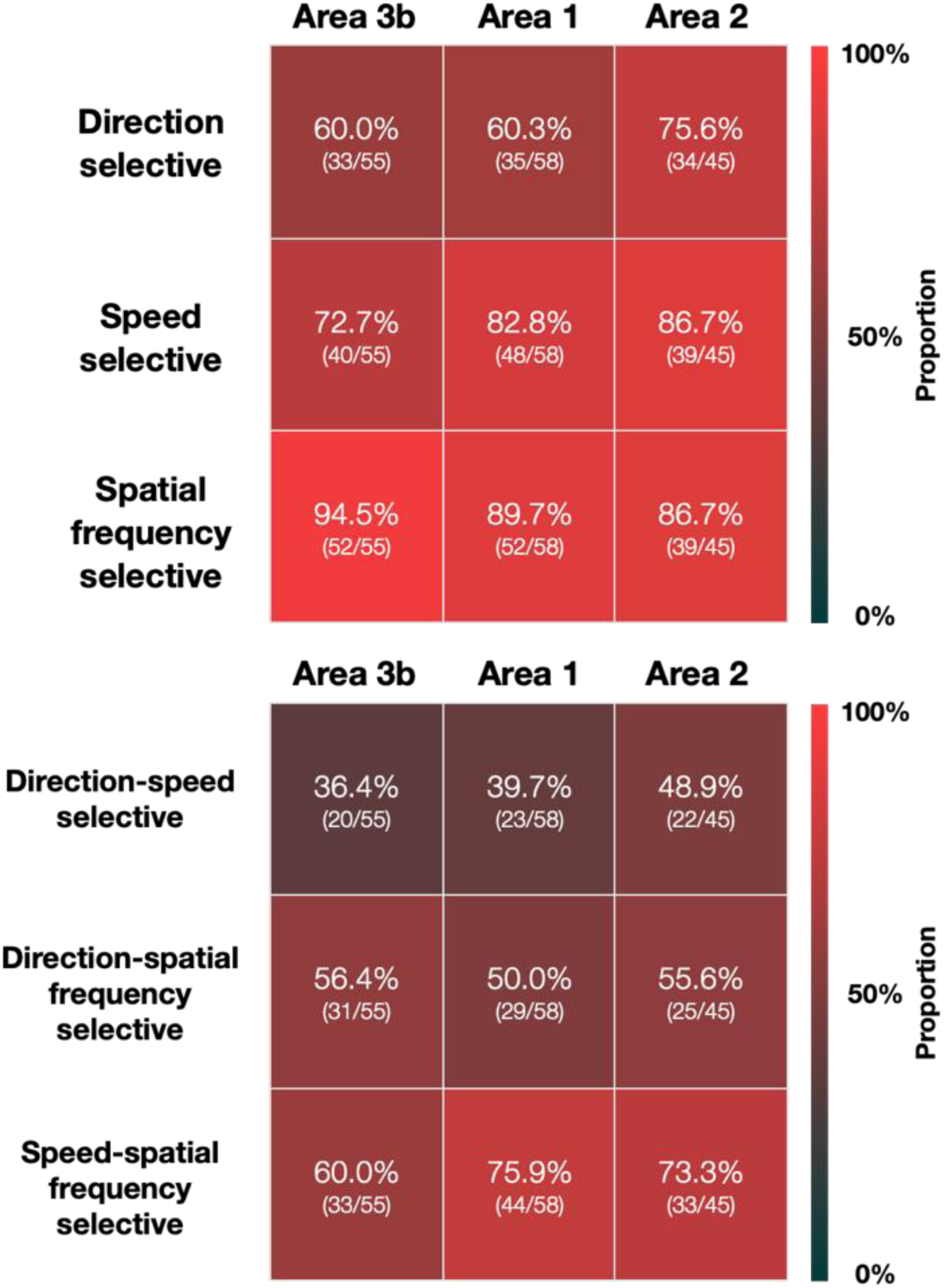
Summary of three-way ANOVA results.

## References

1. Essick, G.K., Franzen, O., and Whitsel, B.L. (1988). Discrimination and scaling of velocity of stimulus motion across the skin. Somatosens Mot Res 6, 21–40. 10.3109/08990228809144639.

2. Bensmaia, S.J., Killebrew, J.H., and Craig, J.C. (2006). Influence of visual motion on tactile motion perception. J Neurophysiol 96, 1625–1637. 10.1152/jn.00192.2006.

3. Dreyer, D.A., Hollins, M., and Whitsel, B.L. (1978). Factors influencing cutaneous directional sensitivity. Sens Processes 2, 71–79.

4. Essick, G.K., Afferica, T., Aldershof, B., Nestor, J., Kelly, D., and Whitsel, B. (1988). Human perioral directional sensitivity. Exp Neurol 100, 506–523. 10.1016/0014-4886(88)90035-0.

5. Norrsell, U., and Olausson, H. (1992). Human, tactile, directional sensibility and its peripheral origins. Acta Physiol Scand 144, 155–161. 10.1111/j.1748-1716.1992.tb09280.x.

6. Gardner, E.P., and Sklar, B.F. (1994). Discrimination of the direction of motion on the human hand: a psychophysical study of stimulation parameters. J Neurophysiol 71, 2414–2429. 10.1152/jn.1994.71.6.2414.

7. Keyson, D.V., and Houtsma, A.J. (1995). Directional sensitivity to a tactile point stimulus moving across the fingerpad. Percept Psychophys 57, 738–744. 10.3758/bf03213278.

8. Delhaye, B.P., O’Donnell, M.K., Lieber, J.D., McLellan, K.R., and Bensmaia, S.J. (2019). Feeling fooled: Texture contaminates the neural code for tactile speed. PLoS Biol 17, e3000431. 10.1371/journal.pbio.3000431.

9. Callier, T., Saal, H.P., Davis-Berg, E.C., and Bensmaia, S.J. (2015). Kinematics of unconstrained tactile texture exploration. J Neurophysiol 113, 3013–3020. 10.1152/jn.00703.2014.

10. Boundy-Singer, Z.M., Saal, H.P., and Bensmaia, S.J. (2017). Speed invariance of tactile texture perception. J Neurophysiol 118, 2371–2377. 10.1152/jn.00161.2017.

11. Handler, A., and Ginty, D.D. (2021). The mechanosensory neurons of touch and their mechanisms of activation. Nature Reviews Neuroscience 22, 521–537.

12. Burton, H., and Fabri, M. (1995). Ipsilateral intracortical connections of physiologically defined cutaneous representations in areas 3b and 1 of macaque monkeys: projections in the vicinity of the central sulcus. Journal of Comparative Neurology 355, 508–538.

13. Martuzzi, R., van der Zwaag, W., Farthouat, J., Gruetter, R., and Blanke, O. (2014). Human finger somatotopy in areas 3b, 1, and 2: a 7T fMRI study using a natural stimulus. Human brain mapping 35, 213–226.

14. Iwamura, Y., Tanaka, M., Sakamoto, M., and Hikosaka, O. (1983). Converging patterns of finger representation and complex response properties of neurons in area 1 of the first somatosensory cortex of the conscious monkey. Experimental Brain Research 51, 327–337.

15. Tremblay, F., Ageranioti-Belanger, S.A., and Chapman, C.E. (1996). Cortical mechanisms underlying tactile discrimination in the monkey. I. Role of primary somatosensory cortex in passive texture discrimination. J Neurophysiol 76, 3382–3403. 10.1152/jn.1996.76.5.3382.

16. Iwamura, Y. (1998). Hierarchical somatosensory processing. Current opinion in neurobiology 8, 522–528.

17. Johnson, K.O., and Phillips, J.R. (1981). Tactile spatial resolution. I. Two-point discrimination, gap detection, grating resolution, and letter recognition. Journal of neurophysiology 46, 1177–1192.

18. Dépeault, A., Meftah el, M., and Chapman, C.E. (2008). Tactile speed scaling: contributions of time and space. J Neurophysiol 99, 1422–1434. 10.1152/jn.01209.2007.

19. Manfredi, L.R., Saal, H.P., Brown, K.J., Zielinski, M.C., Dammann, J.F., 3rd, Polashock, V.S., and Bensmaia, S.J. (2014). Natural scenes in tactile texture. J Neurophysiol 111, 1792–1802. 10.1152/jn.00680.2013.

20. Johnson, K.O., and Phillips, J.R. (1988). A rotating drum stimulator for scanning embossed patterns and textures across the skin. Journal of neuroscience methods 22, 221–231.

21. Pei, Y.C., Lee, T.C., Chang, T.Y., Ruffatto, D., 3rd, Spenko, M., and Bensmaia, S. (2014). A multi-digit tactile motion stimulator. J Neurosci Methods 226, 80–87. 10.1016/j.jneumeth.2014.01.021.

22. Weber, A.I., Saal, H.P., Lieber, J.D., Cheng, J.W., Manfredi, L.R., Dammann, J.F., 3rd, and Bensmaia, S.J. (2013). Spatial and temporal codes mediate the tactile perception of natural textures. Proc Natl Acad Sci U S A 110, 17107–17112. 10.1073/pnas.1305509110.

23. Srinivasan, M.A., and LaMotte, R.H. (1987). Tactile discrimination of shape: responses of slowly and rapidly adapting mechanoreceptive afferents to a step indented into the monkey fingerpad. Journal of Neuroscience 7, 1682–1697.

24. Mountcastle, V.B., LaMotte, R.H., and Carli, G. (1972). Detection thresholds for stimuli in humans and monkeys: comparison with threshold events in mechanoreceptive afferent nerve fibers innervating the monkey hand. Journal of neurophysiology 35, 122–136.

25. Edin, B.B., Essick, G.K., Trulsson, M., and Olsson, K.A. (1995). Receptor encoding of moving tactile stimuli in humans. I. Temporal pattern of discharge of individual low-threshold mechanoreceptors. J Neurosci 15, 830–847. 10.1523/JNEUROSCI.15-01-00830.1995.

26. Essick, G.K., and Edin, B.B. (1995). Receptor encoding of moving tactile stimuli in humans. II. The mean response of individual low-threshold mechanoreceptors to motion across the receptive field. J Neurosci 15, 848–864. 10.1523/JNEUROSCI.15-01-00848.1995.

27. DiCarlo, J.J., and Johnson, K.O. (1999). Velocity invariance of receptive field structure in somatosensory cortical area 3b of the alert monkey. Journal of Neuroscience 19, 401–419.

28. Dépeault, A., Meftah el, M., and Chapman, C.E. (2013). Neuronal correlates of tactile speed in primary somatosensory cortex. J Neurophysiol 110, 1554–1566. 10.1152/jn.00675.2012.

29. Pei, Y.C., Hsiao, S.S., Craig, J.C., and Bensmaia, S.J. (2011). Neural mechanisms of tactile motion integration in somatosensory cortex. Neuron 69, 536–547. 10.1016/j.neuron.2010.12.033.

30. Mountcastle, V.B., Talbot, W.H., Darian-Smith, I., and Kornhuber, H.H. (1967). Neural basis of the sense of flutter-vibration. Science 155, 597–600.

31. Mountcastle, V.B., Talbot, W.H., Sakata, H., and Hyvärinen, J. (1969). Cortical neuronal mechanisms in flutter-vibration studied in unanesthetized monkeys. Neuronal periodicity and frequency discrimination. Journal of neurophysiology 32, 452–484.

32. Gardner, E.P., Palmer, C.I., Hamalainen, H.A., and Warren, S. (1992). Simulation of motion on the skin. V. Effect of stimulus temporal frequency on the representation of moving bar patterns in primary somatosensory cortex of monkeys. J Neurophysiol 67, 37–63. 10.1152/jn.1992.67.1.37.

33. Delhaye, B., Hayward, V., Lefevre, P., and Thonnard, J.L. (2012). Texture-induced vibrations in the forearm during tactile exploration. Front Behav Neurosci 6, 37. 10.3389/fnbeh.2012.00037.

34. Bensmaia, S.J., and Hollins, M. (2003). The vibrations of texture. Somatosens Mot Res 20, 33–43. 10.1080/0899022031000083825.

35. Gamzu, E., and Ahissar, E. (2001). Importance of temporal cues for tactile spatial-frequency discrimination. J Neurosci 21, 7416–7427. 10.1523/JNEUROSCI.21-18-07416.2001.

36. McIntyre, S., Birznieks, I., Vickery, R.M., Holcombe, A.O., and Seizova-Cajic, T. (2016). The tactile motion aftereffect suggests an intensive code for speed in neurons sensitive to both speed and direction of motion. J Neurophysiol 115, 1703–1712. 10.1152/jn.00460.2015.

37. Pei, Y.C., Hsiao, S.S., Craig, J.C., and Bensmaia, S.J. (2010). Shape invariant coding of motion direction in somatosensory cortex. PLoS Biol 8, e1000305. 10.1371/journal.pbio.1000305.

38. Salinas, E., Hernandez, A., Zainos, A., and Romo, R. (2000). Periodicity and firing rate as candidate neural codes for the frequency of vibrotactile stimuli. Journal of neuroscience 20, 5503–5515.

39. Long, K.H., Lieber, J.D., and Bensmaia, S.J. (2022). Texture is encoded in precise temporal spiking patterns in primate somatosensory cortex. Nature communications 13, 1311.

40. Romo, R., Hernández, A., Zainos, A., Lemus, L., and Brody, C.D. (2002). Neuronal correlates of decision-making in secondary somatosensory cortex. Nature neuroscience 5, 1217–1225.

41. Romo, R., and Salinas, E. (2003). Flutter discrimination: neural codes, perception, memory and decision making. Nature Reviews Neuroscience 4, 203–218.

42. Chen, Y.P., Yeh, C.I., Lee, T.C., Huang, J.J., and Pei, Y.C. (2020). Relative posture between head and finger determines perceived tactile direction of motion. Sci Rep 10, 5494. 10.1038/s41598-020-62327-x.

43. Hsu, Y.C., Yeh, C.I., Huang, J.J., Hung, C.H., Hung, C.P., and Pei, Y.C. (2019). Illusory Motion Reversal in Touch. Front Neurosci 13, 605. 10.3389/fnins.2019.00605.

44. Efron, B., and Tibshirani, R. (1986). Bootstrap methods for standard errors, confidence intervals, and other measures of statistical accuracy. Statistical science, 54–75.

45. Feng, W., Havenith, M.N., Wang, P., Singer, W., and Nikolic, D. (2010). Frequencies of gamma/beta oscillations are stably tuned to stimulus properties. Neuroreport 21, 680–684.

46. Pei, Y.-C., and Bensmaia, S.J. (2014). The neural basis of tactile motion perception. Journal of Neurophysiology 112, 3023–3032.

